# Estimating fitness effects of mutations in the presence of genetic linkage

**DOI:** 10.1101/2025.11.26.690712

**Authors:** S.V. Lagunov, I.M. Rouzine

## Abstract

Probabilistic prognosis of the evolution of population requires the knowledge of the fitness effects of mutations at different genomic sites. However, the signature of natural selection is eclipsed by strong noise in data, because the common ancestors of different sites render their evolution inter-dependent. Together with mutation and recombination, genetic linkage also makes evolution stochastic and requires averaging over many independent populations. Here we develop a method designed to work in the presence of strong linkage effects. For testing, we apply it to simulated genomic sequences generated by a Monte Carlo algorithm with known selection coefficients. Results demonstrate the good accuracy of the estimates of relative selection coefficients, if more than ten independent populations are used for averaging. Infrequent recombination partly compensates linkage effects and improves the accuracy. The method is then used to estimate the selection coefficients of 10,000 genomic sites of *E. coli*. These findings enable the inference of adaptive landscape under the conditions of strong multi-site linkage.

**AUTHOR SUMMARY:** Probabilistic prognosis of evolution of an organism requires the knowledge of fitness effects of mutations at different genomic sites. However, genetic linkage between different sites due to their common phylogenetic history obscures the effects of natural selection. Here we develop a linkage-resistant method to infer fitness effects and test its accuracy on mock sequences generated by an evolutionary algorithm with known fitness effects. Results demonstrate fair accuracy when averaging is performed over ten or more independently-evolving populations. After application of this program to genomic data for *E. coli*, relative selection coefficients for thousands of genomic sites are obtained.

## INTRODUCTION

Evolutionary dynamics of a population of DNA sequences is controlled by the interplay of various evolutionary factors, including random mutation, natural selection, genetic drift, and recombination. Of central interest is adaptation to a new environment, which occurs due to emergence of rare mutations that increase the fitness of the organism (average progeny number). Deleterious mutations emerge during adaptation as well. The advantage of each favorable or deleterious mutation is measured by the relative change it causes in fitness. Thus, the knowledge of fitness effects for different mutations is essential for predicting the evolutionary trajectory.

Measuring fitness effects of mutations directly is possible only in some cases, for example, in viruses and bacteria, but even then it requires rather elaborate experiments [1-5]. Therefore, a great effort has been invested in their estimation from genomic data using simplifying models. The average fitness effect of a beneficial mutation in HIV genome was estimated using genetic samples from infected patients and an one-site model [6]. Selection coefficients in the hemagglutinin gene of influenza A H3N2 were estimated by fitting the predictions of one-site and two-site models to the time-series sequence data on the hemagglutinin gene of influenza virus A H3N2 [7]. Another group [8] proposed a method of estimating selection coefficients of deleterious mutations based on an assumption about the form of fitness effect distribution among loci. In SARS-CoV-2, fitness effects of mutations were computed as the natural logarithm of the ratio of their actual to expected counts, with a correction for the codon founder effect [9]. One-locus Wright-Fisher model was used to estimate selection coefficients from genomic data for Sindbis virus [10].

A major complication in predicting the evolutionary trajectory and estimating the fitness effects of mutations is that the evolution of different sites is coupled due to their common phylogeny. Because different sites share ancestors, genetic linkage effects between them arise, including clonal interference and the genetic background effect [11]. Linkage limits the use of one-site and two-site models for the inference of observable properties, and a more complex framework taking into account a large number of genomic sites must be used [12, 13]. The multi-site linkage modifies all evolutionary observables, from the evolution rate and fitness distribution[12, 14] to phylogenetic properties [15-17]. Recombination compensates linkage effects, but only partly [15, 18-20]. The multi–site models have been successfully applied to predict evolutionary parameters of several pathogenic viruses, including HIV [21] and influenza [22-26].

Of great practical importance is the ability to predict virus evolution, in the probabilistic sense, with the goal of optimizing vaccines and antiviral drugs [27], with the caveat that vaccines can expedite the rate of virus evolution [28]. Based on multi-site models, methods of predicting the fitness of whole virus strains from their position on the phylogenetic trees [29] and from the immune memory of extinct strains [24, 30] have been developed. In addition to whole strains, the evolution of specific genomic sites represents biomedical interest as well, which brings us back to estimating their selection coefficients.

A recent modeling work [31] demonstrated that, when many genomic sites adapt slowly, alleles are distributed among genomes and sites to maximize disorder at any one time. The average evolutionary dynamics of separate sites was mapped onto an one-site model, where time is effectively slowed down by a factor, the same for all sites [32, 33]. These findings allowed to develop a method estimating the relative fitness effects of mutations from the evolutionary dynamics, without neglecting linkage effects [33].

In the present work, we test the accuracy of this method [32, 33] on simulated sequences with known fitness effects of mutations. Then, the method is applied to the genomic data on *Escherichia coli* [34].

## RESULTS

### Linkage-proof method to estimate the fitness effect of mutation

We determined the relative influence of individual mutations on fitness (selection coefficient) by using a time series of allelic frequencies at many sites and averaging them across independently-evolving populations. Genomic samples were collected at multiple time points from several independent populations. All genomes were aligned, and the consensus of the entire dataset was found. At each polymorphic site and each time point, the frequency of minority allele was calculated. The relative value of the selection coefficient was estimated for each site, using a procedure described in *Methods*.

### Computer model of population

The accuracy of the method was tested on artificial sequences generated by a Monte Carlo algorithm of Wright–Fisher type. An individual genome was represented by a binary sequence showing an allele at each locus, one of the two possible. In each generation, the population was replaced with a new one, taking into account the effects of mutation, natural selection, random genetic drift, genetic linkage, and occasional recombination. Fitness of a genome (average progeny number) was calculated from the sum of beneficial alleles weighted with their fitness effects set in the program. See *Methods* for the details.

### Testing the accuracy of the method for a maximally–diverse population in the presence of recombination

Before applying the method to real biological data, a series of computational experiments was conducted on simulated binary sequence data to test the accuracy. We applied the algorithm to sequences obtained from simulations, where all parameters are set in advance including the value of selection coefficient, denoted *s*_*i*_, at each genomic site *i*. Using the method described above, for each time point *t*_*m*_, we estimated the relative value of selection coefficient for each site in the form *β*(*t*_*m*_) *s* _*i*_, where *β*(*t*)is a function of time (*Methods*). To verify the accuracy of the predicted values of selection coefficients, they were compared with the known values of selection coefficients. Results demonstrate fairly good linear correlation between the inferred and the actual values (Figure 1A).

**Figure 1.**
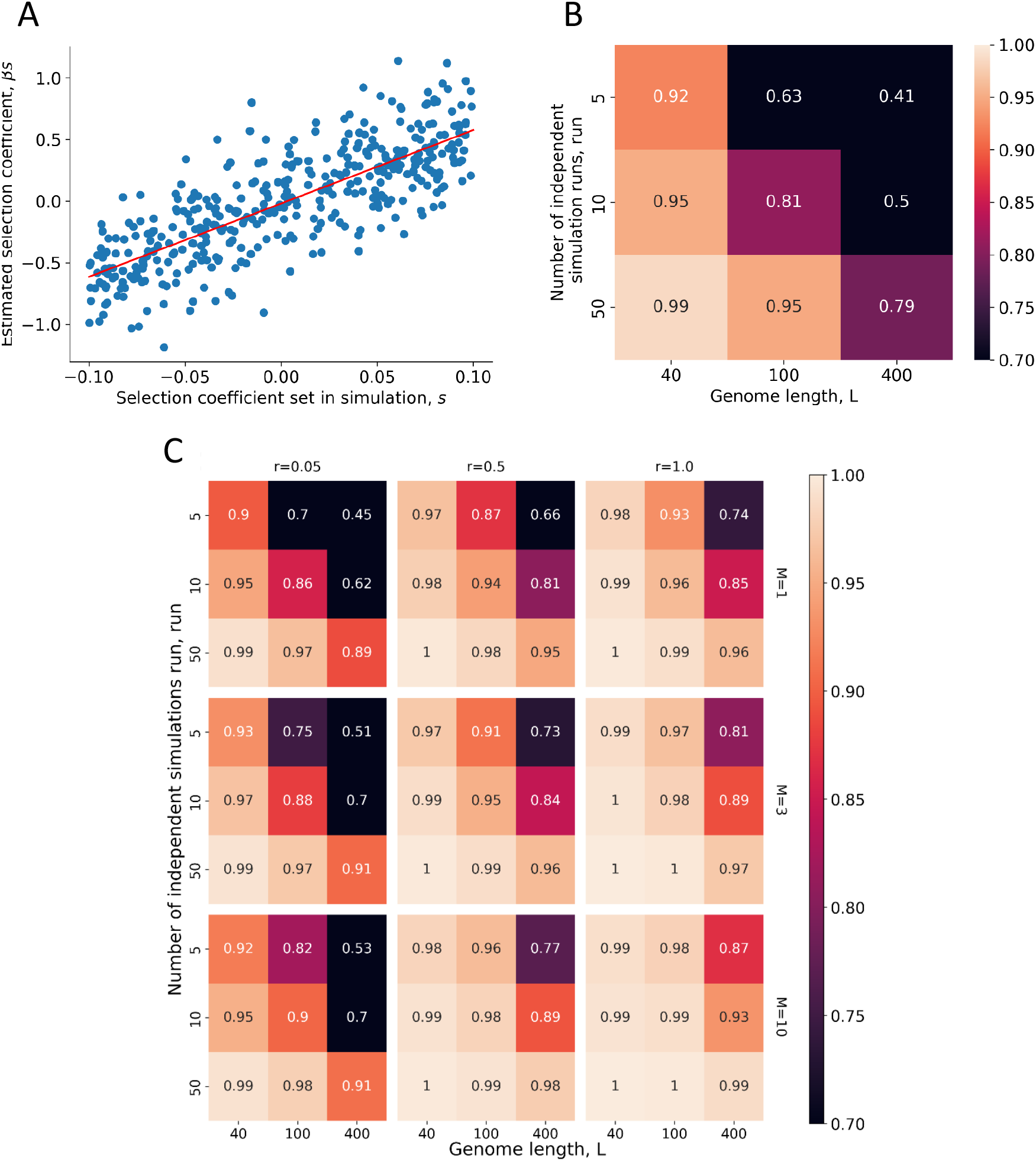
Testing the accuracy of the method for an initial maximally–diverse population. **(A)** X-axis: values of *s* set in the simulation. Y-axis: their estimates in relative units, *βs*. Red line: linear fit with slope *β* = 2.5 and correlation *r*^2^ = 0.46. (For *L* = 100, *r*^2^ = 0.93.) Simulation parameters: initial mutation frequency *f*_0_ = 0.5, mutation rate *µL* = 0.07, uniform distribution of selection coefficients with average *s* = 0.1, population size *N* = 1000, the number of simulation runs *run* = 50, genome length *L* = 400, no recombination. (B, C) Pearson’s correlation coefficient between the actual values of selection coefficients and their estimates at different values of genome length (columns) and numbers of simulation runs (rows). (B) No recombination. (C) Probability of recombination *r* and crossover number *M* are shown. Sensitivity to population size, sample size, and mutation rate is shown in Fig. S1 in Supplement.

To determine under which parameter values the highest accuracy was achieved, Pearson correlation coefficient was calculated between the estimated values of *β*(*t*_*m*_) *s* _*i*_ and the values of *s*_*i*_ set in the simulation. Pearson coefficient was averaged over time points. The results demonstrate that the correlation between the estimated and the actual values decreases with the length of genome due to increasing linkage effects and increases with the number of independent runs and the probability of recombination (Figure 1B and C). This correlation depends weakly on population size, sample size, and mutation rate (Figure S1).

### Testing the method for an initially–maladapted population

We also repeated the same accuracy test for the case of initially poorly adapted population with a 5% of beneficial alleles in the absence of crossover recombination. These conditions are closer to the experiment on bacterial evolution discussed below. The accuracy of the method is still rather high (Fig. 2).

**Figure 2.**
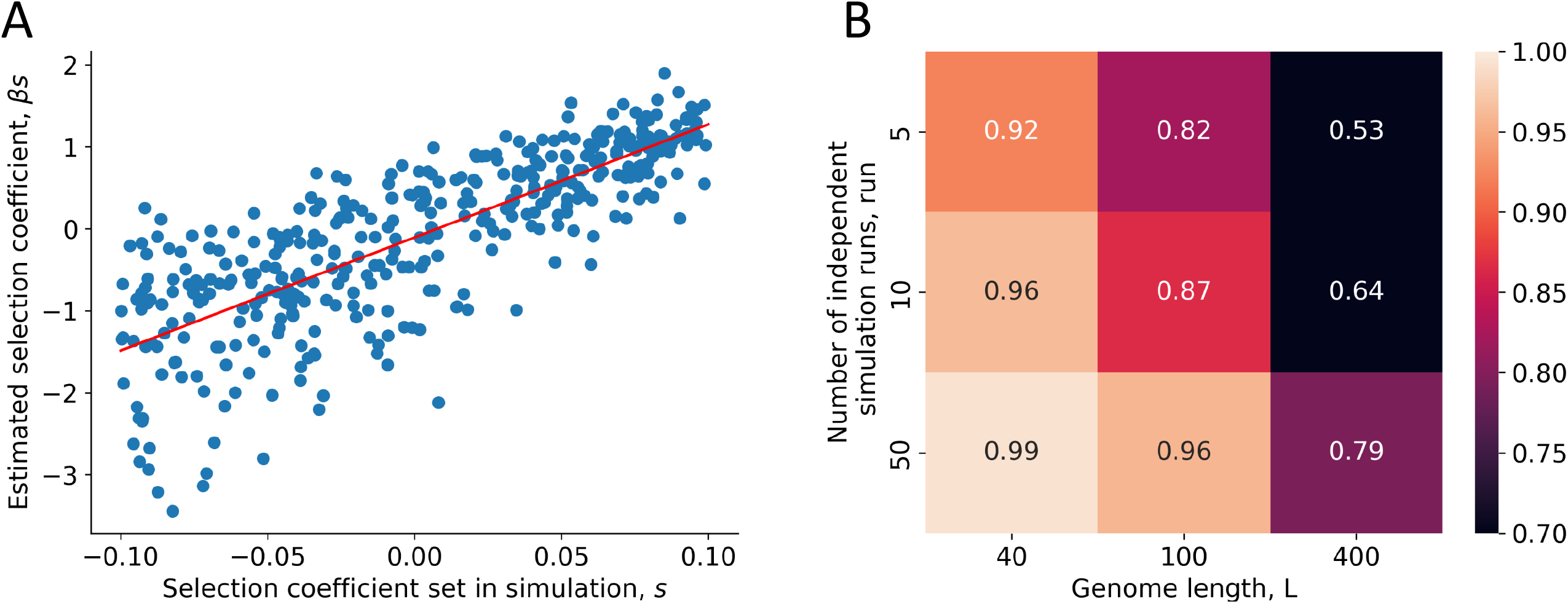
Testing the accuracy of the method for an initial weakly–diverse population (*f*_0_ = 0. 05). Recombination is absent. Notation and other parameters as in Fig. 1.

### Application to genomic data of E. Coli

After validating the method on simulated data, it was applied to data from a real biological population. The sequences were downloaded from a database created in a long-term evolution experiment on *E. coli* [34]. Data analysis was performed for early generations to avoid two issues: the influence of variants with increased mutation probabilities and the population segregation due to differential food use that emerged in several populations over time [34]. The algorithm identified approximately 10,000 evolving sites. The values of relative selection coefficients for some sites are presented in Tables 1 and 2. The distribution of selection coefficients (Figure 3) has a non-monotonous form slightly resembling a distribution previously observed for poliovirus [3]. Note that the number of beneficial mutaions exceeds that of deleterious, because the population in the beginning of adaptation to new nutritions. The full table for all sites is provided in *Supplement* online, file *coeff_s_ecoli*.*csv*.

**Table 1.**
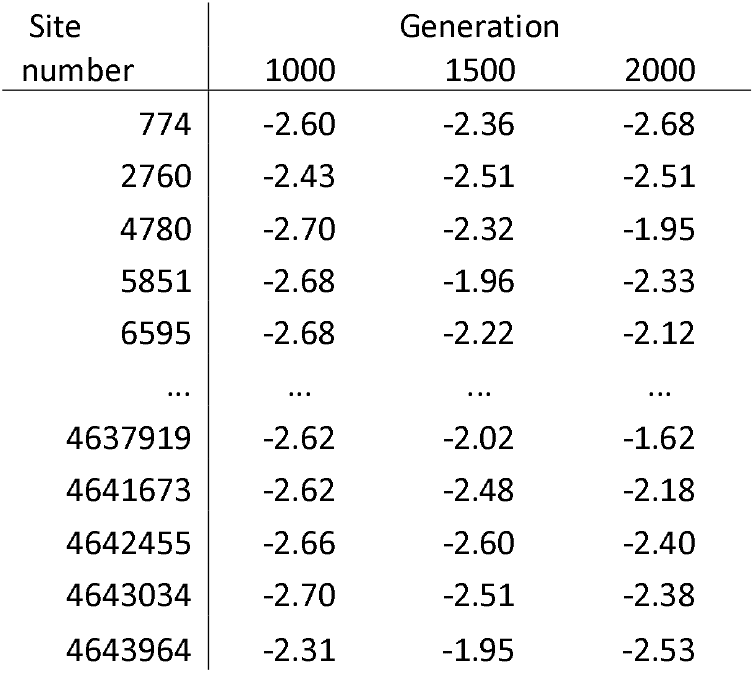
The values of *β*(*t*_*m*_) *s*_*i*_ for three chosen generations of *E. Coli* sorted by site number. Only some genomic sites are shown. The full table for all sites is provided in *Supplement* online in file *coeff_s_ecoli*.*csv*.

| Site<br>number | Generation |  |  |
| --- | --- | --- | --- |
|  | 1000 | 1500 | 2000 |
| 774 | -2.60 | -2.36 | -2.68 |
| 2760 | -2.43 | -2.51 | -2.51 |
| 4780 | -2.70 | -2.32 | -1.95 |
| 5851 | -2.68 | -1.96 | -2.33 |
| 6595 | -2.68 | -2.22 | -2.12 |
| ... | ... | ... | ... |
| 4637919 | -2.62 | -2.02 | -1.62 |
| 4641673 | -2.62 | -2.48 | -2.18 |
| 4642455 | -2.66 | -2.60 | -2.40 |
| 4643034 | -2.70 | -2.51 | -2.38 |
| 4643964 | -2.31 | -1.95 | -2.53 |

**Table 2.** Results from Table 1 sorted by the values of *β* (*t*_*m*_)*s*_*i*_.

| Site<br>number | Generation |  |  |
| --- | --- | --- | --- |
|  | 1000 | 1500 | 2000 |
| 2858601 | -2.70 | -2.66 | -2.72 |
| 3221058 | -2.64 | -2.72 | -2.70 |
| 2078726 | -2.68 | -2.68 | -2.68 |
| 3550522 | -2.70 | -2.60 | -2.72 |
| 329433 | -2.72 | -2.68 | -2.62 |
| ... | ... | ... | ... |
| 3919793 | 2.92 | 2.44 | 3.07 |
| 3918316 | 3.13 | 2.76 | 2.60 |
| 3918337 | 3.05 | 2.98 | 2.56 |
| 3919785 | 3.07 | 2.61 | 2.91 |
| 3919785 | 3.07 | 2.76 | 2.91 |

**Figure 3.**
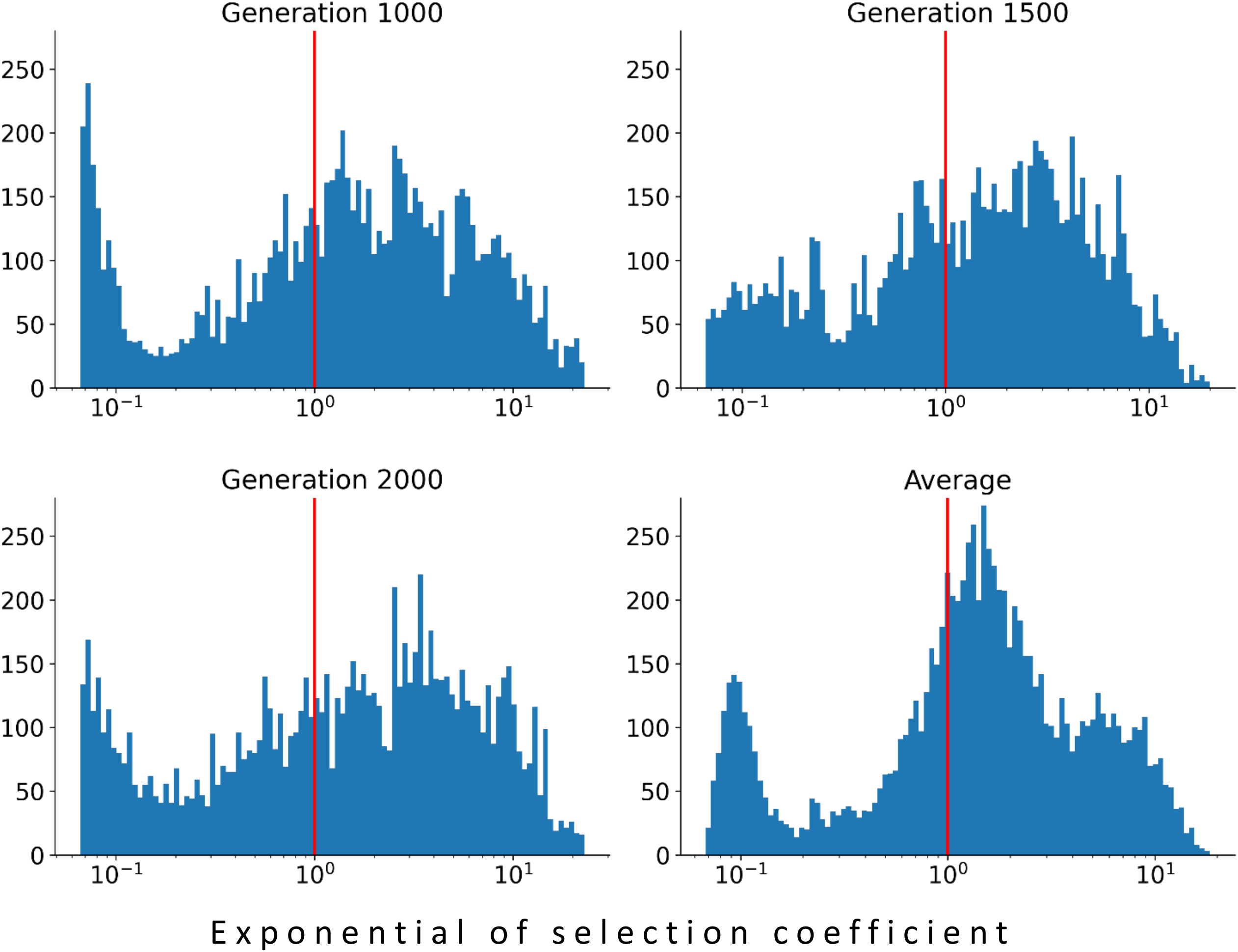
Histogram of the distribution of 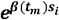 values for E Coli experiment. Values to the right of the red line correspond to positive selection coefficient *s*, and those to the left correspond to negative values. The four panels correspond to three different time points (generations 1000, 1500, and 2000) and the average over time.

## DISCUSSION

The present work offers a software package implementing an algorithm for estimating relative selection coefficients from genomic data despite the presence of strong linkage effects caused by common phylogeny. A test on simulated data demonstrates good accuracy, if ten or more populations are used for averaging. Then the method is used to estimate selection coefficients for 10,000 evolving sites of E. Coli.

The method relies on averaging over multiple independently-evolving populations. This is not a limitation of this specific method of measurement, but a fundamental limitation arising from the stochastic effects of genetic linkage due to the common phylogeny of different sites. Examples of systems where such data can be obtained include various microbiological experiments in multiple serial passages. Another example is the evolution of different metastases in a patient. When addressing human evolution, it is advisable to compare DNA samples of diverged populations from different geographic locations that evolved under similar conditions.

It has been known for a long time that the evolution of a population is strongly affected by the fact that the fates of alleles at different loci are linked unless separated by recombination [11]. Effects of genetic linkage include clonal interference [11, 35], background selection, genetic hitchhiking [36], enhanced accumulation of deleterious mutations (Muller’s ratchet) [12, 37], and the increase of genetic drift at one locus due to selection at another [38]. Genetic linkage also decreases the speed of adaptation due to clonal interference [12, 39]. For this reason, one-locus models usually do not apply in real populations, and multi-locus models based on traveling wave methods must be used [13, 40]. Importantly, genetic linkage makes evolutionary dynamics at different sites stochastic and interdependent, which is a serious obstacle to the inference of fitness landscape from genetic data [41].

To fill this gap, a method based on the approximation of quasi-equilibrium was recently proposed [32, 33]. The cited work demonstrated that the entropy of the distribution of alleles across genomes and sites is maximal conditioned on the value of average population fitness. This approximation, formulated for multi-site adaptation [12], was tested by Monte Carlo simulation [31-33]. The present work is the first application of this method to real genomic data.

After successfully applying to data on *E. Coli*, we also made an attempt to apply this method to data on SARS-CoV-2 and HIV. We found out, however, that the method works only for the systems with a unique best-fit sequence. The method requires averaging allelic frequencies over an ensemble of independently-evolving populations with the same or similar parameters, including the same best-fit sequence. These viruses do not meet this criterion. Indeed, the best-fit sequence for HIV varies between individuals due to the variation of HLA allele subtypes. Coronaviruses, influenza virus and hepatitis C virus are in the state of perpetual antigenic escape from the collective immune memory of population (“Red Queen effect”). In this case, the best-fit sequence does not exist at all. Estimating selection coefficients for such situations will require development of a different approach. The question of generalization of this approach to the system with multiple best-fit sequences remain opens.

Other groups developed a number of methods to estimate selection coefficients neglecting linkage effects. In one work, selection coefficient was estimated from a hidden Markov chain assuming a single-locus model[42]. Another team [43] developed a method for measuring the selection coefficient considering a single locus with two alleles and using data on time series. In their model, backward Kolmogorov equation is used to describe transition probabilities. Another group [7] presented a method for detecting selection signature from population history, by maximizing a likelihood function derived from a two-locus evolutionary model. A likelihood estimation method was used to jointly estimate the selection coefficient and the allele age based on time series data [44]. All these methods neglected linkage effects.

Note that the proportion of beneficial mutations predicted is higher than previously estimated for *E. coli* [45, 46]. The proportion of beneficial alleles is not a universal property of the organism. It varies broadly depending on the state of a population (i.e., time of evolution) and external conditions. Hence the probable reason for discrepancy is that the cited authors propagate *E. Coli*. under different conditions than in [34] whose data we used here. The proportion of deleterious alleles decreases in the process of evolution. If an organism is well–evolved, their deleterious alleles are very rare. Also, the previous method of estimating *s* is based on a 1-locus model.

### Conclusions

Using a method developed to withstand linkage effects, we estimate the relative values of selection coefficient for several thousand sites of the evolving *E. Coli*. genome. These results demonstrate the feasibility of the measurements of selection coefficients from genomic data in populations evolving under directional selection in the presence of strong linkage effects.

## METHODS

### Generating sequences for method testing

Evolution of a haploid asexual population of *N* binary sequences is simulated by a Monte Carlo code. In an individual genome, each locus (site, nucleotide position, amino acid position) numbered *i* = 1,2, …,*L* is occupied by one of two alleles, either the wild-type allele, denoted *K*_*i*_ = 0, or the mutant allele, *K*_*i*_ = 1.We use a discrete generation scheme in the absence of generation overlap (Wright-Fisher model). The evolutionary factors included in the model are random mutation with rate *μL* per genome, constant directional selection, random genetic drift due to random sampling of progeny and recombination with rate *r*. Selection includes an epistatic network with a set strength and topology. The logarithm of the average progeny number of an individual genome, *W*, depends on sequence [*K*_*i*_] as given by

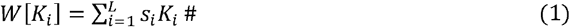

Here selection coefficient *s*_*i*_ denotes the fitness gain from a beneficial mutation at site *i*.

If a population is very poorly adapted to environment, and starts from almost 100% deleterious alleles at every locus, every locus will experience beneficial mutations and, hence, gradually accumulate beneficial alleles, until they are almost fixed in a population [47, 48]. In the middle of this process, population is extremely diverse, and has a 50% of deleterious alleles. Examples include fixation of immune escape and drug–resistant mutations [6, 49, 50]. Thus, the frequency of deleterious alleles varies broadly in time depending how well a population is adapted to environment. If it is well adapted, their frequencies are very low. For sites in the middle of adaption, which was the focus of our study, it is very high.

We consider two adaptation scenarios: (i) initially maximally diverse and (ii) initially poorly adapted population (Figs. 1 and 2). The initial frequency of beneficial alleles is set at *f*_0_ = 0.5, in Fig. 1 and *f*_0_ = 0.05 respectively. The mutation rate *μL* = 0.07, population size *N* = 1000. We have assumed a uniform distribution of selection coefficients with average *s* = 0.1, although other distributions can also be used [51].

In experiment [34] whose data we use below, the authors use poorly adapted E Coli that have a large number of adapting sites. Hence, we considered the scenario case of poorly adapted populations as our main scenario.

Monte-Carlo simulation code is written in MATLAB^TM^ and deposited at https://github.com/Rictograf/s_measurement/tree/main/monte-carlo%20simulation.

### Data processing

We downloaded genomic sequences of *E*.*Coli*. obtained in long-term experiments [34] from a public database https://www.ncbi.nlm.nih.gov/bioproject/?term=PRJNA380528. If sequences contained deletions or unspecified nucleotides (‘-’ or ‘N’), the data were cleaned of them, as follows.

Sequences, in which the number of such occurrences exceeded a certain threshold, were removed. Then sites where the number of deletions and unspecified nucleotides exceeded the *SeqGapsThreshold* threshold were removed. Thus, insertions are not considered by our algorithm, because during alignment they form a column of ‘-’ values at the site where they arise. Next, any remaining ‘-’ or ‘N’ values are considered minor variants and assigned a value of 1.

If the frequency of a variant at a site is below threshold *SiteMonomorphThreshold*, the site is considered monomorphic at that point in time and removed. The result is an *L* × *N* table, *L* where is the total site number, and *N* is the number of genomes considered. To binarize the data, the majority variant averaged over all populations and time points for each site is replaced by 0, and all other (minor) variants at that site are replaced by 1. Thresholds are SeqGapsThreshold = 0.05, SiteGapsThreshold = 0.06, SiteMonomorphThreshold = 0.06.

The sequencing data was mapped to consensus NC_007779 (https://www.ncbi.nlm.nih.gov/nuccore/NC_007779.1/) using the Geneious Prime (https://www.geneious.com/) program. Use tool Map to Reference(s). That program was also used to find allele frequencies *f*_*i*_.

### Estimating selection coefficients from genomic data

To estimate selection coefficients, we employ the method developed in [32, 33]. This method is based on the assumption [31] that the entropy of genomic configurations (distribution of alleles over genomes and sites) is maximal given the fitness, at any one time. The intuitive reason for this assumption, which was confirmed by Monte Carlo simulation [31, 33], is that a slow speed of evolution allows the distribution of alleles across sites and genomes to reach the state statistical quasi-equilibrium at any one time.

The frequency of anti-consensus variants denoted *f*_*i*_ (*t*_*m*_) is calculated for each site *i* and time point *t*_*m*_ = *t*_1_,*t*_2_, …. Next, the relative values of the selection coefficient for each site at each time point are found, as follows. Genomic sites *i* are ordered by the value of *f*_*i*_ averaged over the genome sample and time. The value of *β* (*t*_*m*_)*s*_*i*_ is calculated, for each time *t*_*m*_, as [32, 33]

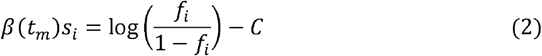

To derive this expression, consider a site with selection coefficient *s* (Eq. 1) in a long genome with total log-fitness W. The models of population genetics (see Discussion) demonstrate that the distribution density of fitness W across individual genomes is a narrow solitary wave moving at a slow speed [13, 40]; in the context of our work, we can consider W fixed. We classify all individual genomes in the population according to the allele at the site: 0 and 1. According to Eq. (1), the contribution of the site to the logarithm of genome fitness depends on the allele: *W*_*site*_ = 0 and *W*_*site*_ = *s*, for alleles 0 and 1, respectively. In the course of evolution, random drift and mutation tend to maximize disorder. On the other hand, the effect of Darwinian selection is to maximize fitness. The standard measure of disorder is configuration entropy *S* defined as the logarithm of the number of possible configurations.

The balance between entropy and fitness is satisfied when entropy is maximal under the restriction that fitness value *W* is fixed (i.e. changes slowly). The maximum value of *S* depends on W, as given by *S* = *S*(*W*). The fitness of the part of genome that does not include the pair is the difference *W* −*W*_*site*_. Hence, entropy of each allele’s subset of sequences is *S* (*W* −*W*_*site*_). Since the genome is long, we can safely assume that *W*_*site*_ is much smaller than *W*, so that the corresponding change in entropy is small, *S* (*W* −*W*_*site*_) ≈ *S* (*W*)− *β* W_*site* ,_ *β* ≡ *ds \ dw* where. The frequency of each haplotype is proportional to the corresponding configuration number, *exp* [*S* (*W* − *W* _*pair*_)], and we arrive at Eq. (2) with *C* = 0. Finally, coefficient *C* is introduced to account for the difference between the best fit sequence and the consensus sequence in experiment.

In Eq. 2, the initial value of normalization factor *C* is set to 0 (Fig. 4, bottom). Then, *C* is determined as the ordinate of the intersection point of all curves obtained for different time points *t*_*m*_. To find this point, each curve is approximated by a fourth-degree polynomial (Fig. 4). Thus, we obtain the estimate of the selection coefficients of all sites multiplied by the common coefficient *β* (*t*_*m*_), which depends on time.

**Figure 4.**
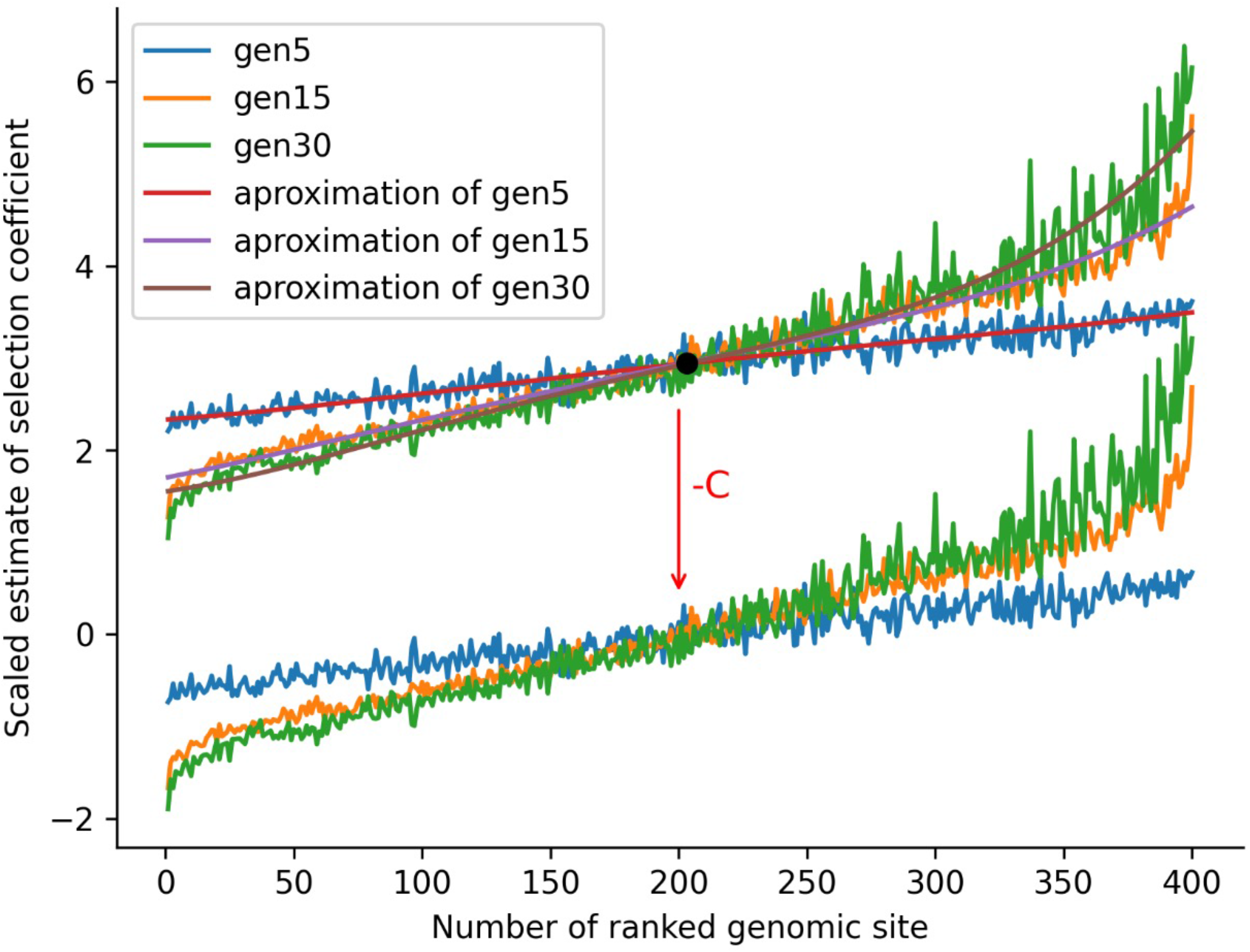
Ranked curves from Eq. 2 with and without C for three time points for simulated sequences. Algorithm visualization. *f*_0_ = 0.05 is the initial mutation rate, mutation rate per genome *µL* = 0.07, average selection coefficient *s* = 0.1, *N* = 1000 individuals in the population, *r* = 0 is the recombination probability, *run* = 50 is the number of simulation runs, *L* = 400 is the genome length. Time points in generations are shown in the legend.

Above procedure is illustrated, for a representative parameter set, in Figures 1 and 2, using simulated sequences produced by Monte Carlo simulation. To test its accuracy, the estimated values of selection coefficients are plotted against their values set in simulation that generated genomic sequences, and Pearson correlation coefficients are calculated.

## Supporting information

Response to Reviewer

## Acknowledgements

This work originated in long discussions with Igor Likhachev. We thank A.A. Makashov for technical help and discussion. We are grateful to A. I. Pikhulya, D. A. Gorshkova, E. A. Vigovskaya, and E. I. Stroganova for technical help.

## Ethical statement

No ethical statement is required.

## Data and software

No new data have been generated in the present work. The program codes are available at https://github.com/Rictograf/s_measurement.

## Supplemental information

Figure S1. Correlations between the estimated and the actual values of selection coefficient: sensitivity to population size (X axis), sample size, and mutation rate (Y axis). The version of Figure 2 with other variable parameters. Fixed parameters: initial allele frequency *f* _0_ = 0.5, average selection coefficient *s* = 0.1, no recombination, the number of simulation runs *run* = 10, genome length *L* = 100.

Supplemental table https://github.com/Rictograf/s_measurement/blob/main/coeff_s_ecoli.csv

## REFERENCES

1. Imhof M, Schlotterer C. Fitness effects of advantageous mutations in evolving Escherichia coli populations. Proc Natl Acad Sci U S A. 2001;98(3):1113–7. doi: 10.1073/pnas.98.3.1113. PubMed PMID: 11158603; PubMed Central PMCID: PMCPMC14717.

2. Kassen R, Bataillon T. Distribution of fitness effects among beneficial mutations before selection in experimental populations of bacteria. Nat Genet. 2006;38(4):484–8. doi: 10.1038/ng1751. PubMed PMID: 16550173.

3. Acevedo A, Brodsky L, Andino R. Mutational and fitness landscapes of an RNA virus revealed through population sequencing. Nature. 2014;505(7485):686–90. Epub 2013/11/29. doi: 10.1038/nature12861. PubMed PMID: 24284629; PubMed Central PMCID: PMC4111796.

4. Stern A, Bianco S, Yeh MT, Wright C, Butcher K, Tang C, et al. Costs and benefits of mutational robustness in RNA viruses. Cell Rep. 2014;8(4):1026–36. doi: 10.1016/j.celrep.2014.07.011. PubMed PMID: 25127138; PubMed Central PMCID: PMCPMC4142091.

5. Wrenbeck EE, Azouz LR, Whitehead TA. Single-mutation fitness landscapes for an enzyme on multiple substrates reveal specificity is globally encoded. Nat Commun. 2017;8:15695. doi: 10.1038/ncomms15695. PubMed PMID: 28585537; PubMed Central PMCID: PMCPMC5467163.

6. Rouzine IM, Coffin JM. Search for the mechanism of genetic variation in the pro gene of human immunodeficiency virus. J Virol. 1999;73(10):8167–78. Epub 1999/09/11. doi: 10.1128/JVI.73.10.8167-8178.1999. PubMed PMID: 10482567; PubMed Central PMCID: PMCPMC112834.

7. Illingworth CJ, Mustonen V. Distinguishing driver and passenger mutations in an evolutionary history categorized by interference. Genetics. 2011;189(3):989–1000. Epub 20110906. doi: 10.1534/genetics.111.133975. PubMed PMID: 21900272; PubMed Central PMCID: PMCPMC3213378.

8. Keightley PD, Eyre-Walker A. Joint inference of the distribution of fitness effects of deleterious mutations and population demography based on nucleotide polymorphism frequencies. Genetics. 2007;177(4):2251–61. doi: 10.1534/genetics.107.080663. PubMed PMID: 18073430; PubMed Central PMCID: PMCPMC2219502.

9. Bloom JD, Neher RA. Fitness effects of mutations to SARS-CoV-2 proteins. Virus Evol. 2023;9(2):vead055. Epub 20230918. doi: 10.1093/ve/vead055. PubMed PMID: 37727875; PubMed Central PMCID: PMCPMC10506532.

10. Silva JMF, Olmo-Uceda MJ, Morley VJ, Turner PE, Elena SF. Taylor’s Power Law rules the dynamics of allele frequencies during viral evolution in response to host changes. J R Soc Interface. 2025;22(228):20250146. Epub 20250716. doi: 10.1098/rsif.2025.0146. PubMed PMID: 40664232; PubMed Central PMCID: PMCPMC12303090.

11. Fisher RA. The genetical theory of natural selection. Oxford, United Kingdom: Clarendon Press, 1958; 1930.

12. Rouzine IM, Wakeley J, Coffin JM. The solitary wave of asexual evolution. Proc Natl Acad Sci U S A. 2003;100(2):587–92. Epub 2003/01/15. doi: 10.1073/pnas.242719299. PubMed PMID: 12525686; PubMed Central PMCID: PMCPMC141040.

13. Rouzine IM. Mathematical modeling of evolution. Volume 1: one-locus and multi-locus theory and recombination De Gruyter, Berlin/Boston; 2020.

14. Desai MM, Fisher DS. Beneficial mutation selection balance and the effect of linkage on positive selection. Genetics. 2007;176(3):1759–98. Epub 2007/05/08. doi: 10.1534/genetics.106.067678. PubMed PMID: 17483432; PubMed Central PMCID: PMCPMC1931526.

15. Rouzine IM, Coffin JM. Highly fit ancestors of a partly sexual haploid population. Theor Popul Biol. 2007;71(2):239–50. Epub 2006/11/14. doi: 10.1016/j.tpb.2006.09.002. PubMed PMID: 17097121; PubMed Central PMCID: PMCPMC1994660.

16. Desai MM, Walczak AM, Fisher DS. Genetic diversity and the structure of genealogies in rapidly adapting populations. Genetics. 2013;193(2):565–85. Epub 2012/12/12. doi: 10.1534/genetics.112.147157. PubMed PMID: 23222656; PubMed Central PMCID: PMCPMC3567745.

17. Neher RA, Hallatschek O. Genealogies of rapidly adapting populations. Proc Natl Acad Sci U S A. 2013;110(2):437–42. Epub 2012/12/28. doi: 10.1073/pnas.1213113110. PubMed PMID: 23269838; PubMed Central PMCID: PMCPMC3545819.

18. Rouzine IM, Coffin JM. Evolution of human immunodeficiency virus under selection and weak recombination. Genetics. 2005;170(1):7–18. Epub 2005/03/04. doi: 10.1534/genetics.104.029926. PubMed PMID: 15744057; PubMed Central PMCID: PMCPMC1449738.

19. Neher RA, Shraiman BI, Fisher DS. Rate of adaptation in large sexual populations. Genetics. 2010;184(2):467–81. Epub 2009/12/02. doi: 10.1534/genetics.109.109009. PubMed PMID: 19948891; PubMed Central PMCID: PMCPMC2828726.

20. Rouzine IM. Long-range linkage effects in adapting sexual populations. Sci Rep. 2023;13(1):12492. Epub 20230801. doi: 10.1038/s41598-023-39392-z. PubMed PMID: 37528175; PubMed Central PMCID: PMCPMC10393966.

21. Rouzine IM, Weinberger LS. The quantitative theory of within-host viral evolution. J Stat Mech: Theory and Experiment. 2013;2013. doi: 10.1088/1742-5468/2013/01/P01009.

22. Lin J, Andreasen V, Casagrandi R, Levin SA. Traveling waves in a model of influenza A drift. J Theor Biol. 2003;222(4):437–45. Epub 2003/06/05. doi: 10.1016/s0022-5193(03)00056-0. PubMed PMID: 12781742.

23. Bedford T, Riley S, Barr IG, Broor S, Chadha M, Cox NJ, et al. Global circulation patterns of seasonal influenza viruses vary with antigenic drift. Nature. 2015;523(7559):217–20. Epub 2015/06/09. doi: 10.1038/nature14460. PubMed PMID: 26053121; PubMed Central PMCID: PMCPMC4499780.

24. Rouzine IM, Rozhnova G. Antigenic evolution of viruses in host populations. PLoS Pathog. 2018;14(9):e1007291. Epub 2018/09/13. doi: 10.1371/journal.ppat.1007291. PubMed PMID: 30208108; PubMed Central PMCID: PMCPMC6173453.

25. Yan L, Neher RA, Shraiman BI. Phylodynamic theory of persistence, extinction and speciation of rapidly adapting pathogens. Elife. 2019;8. Epub 2019/09/19. doi: 10.7554/eLife.44205. PubMed PMID: 31532393; PubMed Central PMCID: PMCPMC6809594.

26. Marchi J, Lassig M, Walczak AM, Mora T. Antigenic waves of virus-immune coevolution. Proc Natl Acad Sci U S A. 2021;118(27). Epub 2021/06/30. doi: 10.1073/pnas.2103398118. PubMed PMID: 34183397; PubMed Central PMCID: PMCPMC8271616.

27. Morris DH, Gostic KM, Pompei S, Bedford T, Luksza M, Neher RA, et al. Predictive Modeling of Influenza Shows the Promise of Applied Evolutionary Biology. Trends Microbiol. 2018;26(2):102–18. Epub 20171030. doi: 10.1016/j.tim.2017.09.004. PubMed PMID: 29097090; PubMed Central PMCID: PMCPMC5830126.

28. Rouzine IM, Rozhnova G. Evolutionary implications of SARS-CoV-2 vaccination for the future design of vaccination strategies. Commun Med (Lond). 2023;3(1):86. Epub 20230619. doi: 10.1038/s43856-023-00320-x. PubMed PMID: 37336956; PubMed Central PMCID: PMCPMC10279745.

29. Neher RA, Russell CA, Shraiman BI. Predicting evolution from the shape of genealogical trees. Elife. 2014;3. Epub 2014/11/12. doi: 10.7554/eLife.03568. PubMed PMID: 25385532; PubMed Central PMCID: PMCPMC4227306.

30. Luksza M, Lassig M. A predictive fitness model for influenza. Nature. 2014;507(7490):57–61. Epub 2014/02/28. doi: 10.1038/nature13087. PubMed PMID: 24572367.

31. Pedruzzi G, Barlukova A, Rouzine IM. Evolutionary footprint of epistasis. PLoS Comput Biol. 2018;14(9):e1006426. Epub 2018/09/18. doi: 10.1371/journal.pcbi.1006426. PubMed PMID: 30222748; PubMed Central PMCID: PMCPMC6177197.

32. Barlukova A, Rouzine IM. The evolutionary origin of the universal distribution of mutation fitness effect. PLoS Comput Biol. 2021;17(3):e1008822. Epub 2021/03/09. doi: 10.1371/journal.pcbi.1008822. PubMed PMID: 33684109; PubMed Central PMCID: PMCPMC7971868.

33. Likhachev IV, Rouzine IM. Measurement of selection coefficients from genomic samples of adapting populations by computer modeling. STAR Protoc. 2023;4(1):101821. Epub 20230304. doi: 10.1016/j.xpro.2022.101821. PubMed PMID: 36871222; PubMed Central PMCID: PMCPMC9999197.

34. Good BH, McDonald MJ, Barrick JE, Lenski RE, Desai MM. The dynamics of molecular evolution over 60,000 generations. Nature. 2017;551(7678):45–50. Epub 20171018. doi: 10.1038/nature24287. PubMed PMID: 29045390; PubMed Central PMCID: PMCPMC5788700.

35. Muller HJ. Some genetic aspects of sex. Am Nat. 1932; 66:118–28.

36. Rice SH. Evolutionary Theory: Mathematical And Conceptual Foundations: Sinauer Associated; 2004.

37. Felsenstein J. The evolutionary advantage of recombination. Genetics. 1974;78(2):737–56. doi: 10.1093/genetics/78.2.737. PubMed PMID: 4448362; PubMed Central PMCID: PMCPMC1213231.

38. Hill WG, Robertson A. The effect of linkage on limits to artificial selection. Genet Res. 1966;8(3):269–94. PubMed PMID: 5980116.

39. Gerrish PJ, Lenski RE. The fate of competing beneficial mutations in an asexual population. Genetica. 1998;102-103(1-6):127–44. Epub 1998/08/28. PubMed PMID: 9720276.

40. Rouzine IM. Mathematical Models of Evolution. Volume 2: Fitness Landscape, Red Queen, Evolutionary Enigmas, and Applications to Virology. Berlin/Boston: De Gruyter; 2023. 305 p.

41. Wei W-H, Hemani G, Haley CS. Detecting epistasis in human complex traits. Nature Reviews Genetics. 2014;15:722.

42. Mathieson I, McVean G. Estimating selection coefficients in spatially structured populations from time series data of allele frequencies. Genetics. 2013;193(3):973–84. Epub 20130110. doi: 10.1534/genetics.112.147611. PubMed PMID: 23307902; PubMed Central PMCID: PMCPMC3584010.

43. Bollback JP, York TL, Nielsen R. Estimation of 2Nes from temporal allele frequency data. Genetics. 2008;179(1):497–502. doi: 10.1534/genetics.107.085019. PubMed PMID: 18493066; PubMed Central PMCID: PMCPMC2390626.

44. Malaspinas AS, Malaspinas O, Evans SN, Slatkin M. Estimating allele age and selection coefficient from time-serial data. Genetics. 2012;192(2):599–607. Epub 20120730. doi: 10.1534/genetics.112.140939. PubMed PMID: 22851647; PubMed Central PMCID: PMCPMC3454883.

45. Elena SF, Ekunwe L, Hajela N, Oden SA, Lenski RE. Distribution of fitness effects caused by random insertion mutations in Escherichia coli. Genetica. 1998;102-103(1-6):349–58. PubMed PMID: 9720287.

46. Eyre-Walker A, Keightley PD. The distribution of fitness effects of new mutations. Nat Rev Genet. 2007;8(8):610–8. doi: 10.1038/nrg2146. PubMed PMID: 17637733.

47. Nielsen R, Slatkin M. Introduction to population genetics: theory and applications: Sinauer Associates; 2013.

48. Rouzine IM, Rodrigo A, Coffin JM. Transition between stochastic evolution and deterministic evolution in the presence of selection: general theory and application to virology [review]. Microbiol Mol Biol Rev. 2001;65(1):151–85.

49. Batorsky R, Kearney MF, Palmer SE, Maldarelli F, Rouzine IM, Coffin JM. Estimate of effective recombination rate and average selection coefficient for HIV in chronic infection. Proc Natl Acad Sci U S A. 2011;108(14):5661–6. Epub 2011/03/26. doi: 10.1073/pnas.1102036108. PubMed PMID: 21436045; PubMed Central PMCID: PMCPMC3078368.

50. Rouzine IM, Weinberger LS. Design requirements for interfering particles to maintain coadaptive stability with HIV-1. J Virol. 2013;87(4):2081–93. Epub 2012/12/12. doi: 10.1128/JVI.02741-12. PubMed PMID: 23221552; PubMed Central PMCID: PMCPMC3571494.

51. Gower G, Pope NS, Rodrigues MF, Tittes S, Tran LN, Alam O, et al. Accessible, Realistic Genome Simulation with Selection Using stdpopsim. Mol Biol Evol. 2025;42(11). doi: 10.1093/molbev/msaf236. PubMed PMID: 40994024; PubMed Central PMCID: PMCPMC12574676.

