## Supplementary material for "Estimating fitness effects of mutations in the presence of genetic linkage": Response to Reviewer

### Response to Reviewer 3

**I have read the response to my comments from the authors. I have previously mentioned that there were three major problems that would be good to address by the authors. I think these three problems were not solved by the authors and I will explain more here below.**

We thank Reviewer for the long time spent on reading our manuscript and response.

From all the Reviewer's comments, we agree with the comment on checking robustness of the method with respect to the initial frequency of deleterious alleles. To address it, we conducted additional simulation for a poorly adapted population with very few initial benefical alleles (Fig. 2). We also carefully repeated our simulations, regrouped figures for better viewing, revised Methods for transparency, and made numerous small edits in the manuscript.

Otherwise, as demonstrated below, "major problems" claimed in Reviewer's report are imaginary and caused by an obvious mismatch between Reviewer's background and the scientific area of our work. While most of the text is written for the general readership, Methods is the only section that requires an expert in mathematical biology, specifically, in evolutionary modeling and mathematical population genetics, as opposed to a biostatistician, bioinformatician, or a wetlab scientist.

The same limitation applies to all multidisciplinary papers. Science has become very specialized. We would never offer critique of the methods of a paper in the field of bioinformatics or experimental biology, as that would be quite unprofessional.

Note that the senior author of the present study, I.M.R., worked for 30 years at three virological labs in USA, Sorbonne in France, and two institutions of the Russian Academy of Sciences, resulting in 45 articles on mathematical biology in Cell, Cell Host and Microbe, PNAS, PLoS Comp Bio and PLoS Pathogens, J.Virology. He trained many students and postdocs, taught at UCLA, USF, Sorbonne, and reviewed for 40+ journals. Previously, he worked in theoretical physics (30 publications). For his full CV, see https://www.iephb.ru/en/employes/rouzine-i/.

**The three problems to solve were: 1) Presentation of the methods. The Reviewer #2 and I have problems understanding the approach of the paper. The authors have added some text that does not help to clarify the methodology of the paper. ... Broadly, I could not follow the methodology and, therefore, the first major problem I mentioned is not solved.**

The fact that Reviewer, not being an expert in the area of evolutionary modeling (see his or her statements below), cannot follow our methodology, is to be expected and quite normal. It is not a "major problem", but a basic fact of life. Definitely, it is not a problem with the manuscript. We all know only what we study for a long time. The major problem is that Reviewer fails to recognize that the manuscript is not in his or her field.

**The authors added the following text: Consider a site with selection coefficient − (Eq. 1) in a long genome with total log-fitness W. The models of population genetics (see Discussion) demonstrate that the distribution density of fitness W across individual genomes is a narrow solitary wave moving at a slow speed; in the context of our work, we can consider W fixed. We classify all individual genomes in the population according to the allele at the site: 0 and 1. According to Eq. (1), the contribution of the site to the logarithm of genome fitness depends on the allele: = 0 and = − , for alleles 0 and 1, respectively. In the course of evolution, random drift and mutation tend to maximize disorder. On the other hand, the effect of Darwinian selection is to maximize fitness. The standard measure of disorder is configuration entropy defined as the logarithm of the number of possible configurations. The balance between entropy and fitness is satisfied when entropy is maximal under the restriction that fitness value is fixed (i.e. changes slowly). The maximum value of depends on W, as given by = ( ). The fitness of the part of genome that does not include the pair is the difference − . Hence, entropy of each allele's subset of sequences is ( − ). Since the genome is long, we can safely assume that is much smaller than , so that the corresponding change in entropy is small, ( − ) ≈ ( ) − , where ≡ / . The frequency of each haplotype is proportional to the corresponding configuration number, [ ( − )], and we arrive at Eq. (2) with = 0. Finally, coefficient is introduced to account for the difference between the best fit sequence and the consensus sequence in experiment......There are assumptions which are not backed up. The authors state that "The models of population genetics (see Discussion) demonstrate that the distribution density of fitness W across individual**

**genomes is a narrow solitary wave moving at a slow speed;" which is not explained or backed up by a reference. I do not comprehend the assumption.**

This is not an "assumption", but a famous result of the broad class of multi-locus models. Traveling wave was also observed experimentally by Desai group. The references are given in Discussion and now copied to this passage. For education, see the book by Rouzine 2020, "Mathematical Modeling of Evolution", De Gruyter.

**"Hence, entropy of each allele's subset of sequences is ( − )." I am unsure what does this mean.**

It was defined above: "The standard measure of disorder is configuration entropy defined as the logarithm of the number of possible configurations

**"The frequency of each haplotype is proportional to the corresponding configuration number, [ ( − )]" I am not sure what this means.**

It refers to the standard definition of mathematical probability of an event as the number of the suitable configurations divided by the number of all possible configurations.

**2) I also mentioned a second problem: "The second problem is the simulations presented to validate the methods. The authors use a simulation scheme where the initial frequency of deleterious alleles, f0 = 0.5. Deleterious alleles tend to have low frequencies (Nielsen et al., 2005) making this simulation scheme not particularly realistic." The authors mention that: "Frequency of deleterious alleles vary broadly depending on a locus and how well a population is adapted to environment. If it is well adapted, their frequencies are very low. For sites in the middle of adaption, which was the focus of our study, it is very high. In the cited experiment we fit, the authors use poorly adapted E Coli to have a large number of adapting sites. Hence, we used the case of poorly adapted populations for testing." The statement "For sites in the middle of adaption, which was the focus of our study, it is very high." Is not backed up by any reference and is not true.**

It is true. Moreover, it is common knowledge in population genetics that loci in the middle of adaptation are highly diverse. During the process adaptation, the frequency of beneficial alleles gradually passes from almost 0 to almost 100. In the revised version, we added citations [47, 48] to the textbook on basic population genetics by Nielsen and Slatkin and a popular review on the same by Rouzine et al 2001. We also clarify in revision:

If a population is very poorly adapted to environment, and starts from 100% deleterious alleles at every locus, every locus will then gradually accumulate beneficial alleles due to positive Darwinian selection, until they are almost fixed in a population [47, 48]. In the middle of this process, population is extremely diverse, and has a 50% of deleterious alleles. Examples include fixation of immune escape and drug resistant mutations [6, 49, 50]. Thus, the frequency of deleterious alleles varies broadly depending on a locus and how well a population is adapted to environment. If it is well adapted, their frequencies are very low. For sites in the middle of adaption, which was the focus of our study, it is very high. In the cited experiment we fit, the authors use poorly adapted E Coli to have a large number of adapting sites. Hence, we used the case of poorly adapted populations for testing.

**In the reference I mentioned to the authors (Nielsen et al., 2005), the authors could see that alleles at a frequency of 0.5 are rare under negative selection or neutrality and at low frequency under positive selection (Figure 2).**

Reviewer compares apples with oranges. These are two different biological systems. Richard Lensky's team started the long-term adaptation experiment by adding new nutritions to which bacteria had to adapt for thousands of generations. Nielsen et al considered a system close to equilibrium. Of course, allelic frequencies, which also depend on time, are very different between the two systems.

**Even if the alleles are "in the middle of adaption", the alleles would not tend to be at a frequency of 0.5, and the authors do not provide any good argument to explore this case.**

Yes, they would be at 50%, and this fact is common knowledge in population genetics. In the middle of this process, many sites will be diverse and have 50% of each allele (see for example, the model and data on the evolution of HIV in Rouzine and Coffin J. Virol. 1999).

**If the E. Coli are poorly adapted, there is no obvious reason for the alleles to have a frequency of 0.5. Therefore the authors are testing an unrealistic simulation scenario...**

The only comment in the Report we partly agree with. Although such scenario is not unrealistic, it does not describe the start of Lensky's experiment we use. To address this concern, we repeated our simulation for poorly adapted population, with the initial beneficial allelic frequency 0.05 (see new Fig. 2).

**The authors state: "We note that the cited paper does not test other teams' methods either." and that is precisely what is done in Figure 3. Therefore, this statement is false.**
**The authors state: "We note that the cited paper does not test other teams' methods either. Nobody really does it in Articles. If we are ever invited to write a Review on this topic, we will be more than happy to compare various approaches.". That is a false statement. It is very normal to compare a method to other methods that try to solve the same problem. The Gower paper compares different methods to estimate the DFE. And many papers compare their method to other methods ( https://journals.plos.org/plosgenetics/article?id=10.1371/journal.pgen.1008384 , https://academic.oup.com/mbe/article/39/1/msab332/6433161?guestAccessKey= ).**

Some authors compare their method with other methods**, when such comparison makes sense.** This is not the case here. As stated in Introduction and Abstract, the aim is to propose a new method of estimating s and to test its accuracy by the means of a numerical control experiment, where artificial genome sequences are generated in Monte Carlo simulation with known s. Comparison with other methods, whose own accuracy was not tested in a similar fashion, is not informative.

Moreover, comparison of the results of a technique based on a multilocus to them is apples and oranges, because it is designed for a different regime, when the total number of mutations in a population per generation is very large.

**My request for the authors was that "If possible, it would be good to verify the results doing simulations using the simulator SLiM, setting a realistic distribution of fitness effects for the deleterious mutations and then adding mutations under a particular selection coefficient to be estimated with their method. Particularly it would be good to test their method using realistic simulations (see Gower et al., 2025 Molecular Biology and Evolution)." The authors refused to do that because: "Also, adding more details to a model does not necessarily makes it more "realistic" (or accurate). On the contrary, a more complex model usually accumulates errors, if the key factors are not well-controlled. Unfortunately, Gower et al do not describe their population genetics model in the detail in Methods for us to comment on it, in particular, on the critical inclusion of genetic linkage."...**
**...Testing the robustness of the methods to different assumptions is important. The authors do not need to "add details to a model". They just need to show how robust is their method to "factors that are not well-controlled" but could be affecting the inference. That is why I suggested to do SLiM simulations.**
**The authors state that "Unfortunately, Gower et al do not describe their population genetics model in the detail in Methods for us to comment on it, in particular, on the critical inclusion of genetic linkage.". The authors can read the extensive documention from Gower et al. 2005 ( https://popsim-consortium.github.io/stdpopsim-docs/stable/index.html ) to learn how to do these simulations and how genetic linkage is included. Otherwise, testing their method under reasonable SLiM simulations would be very helpful to test that the method from the authors provides reasonable inferences.**

The fact that Reviewer religiously believes SLIM "realistic and reasonable", but cannot explain how it works and refers to "extensive documentation", is very telling.

Our aim is to test our method on simulated sequences for an example of a broadly distributed selection coefficient. We do not need external software for this aim. Our team of mathematical biologists writes custom-made programs for the mathematical models we develop and analyze ourselves for 30 years. Using external programs with unknown content is not "testing robustness of the method of estimating s". It is "testing somebody else's software ". This is not our job.

**3) My third problem is that "The third problem from this paper is that the authors find a high proportion of advantageous mutations with their method which is very different from what has been previously estimated on E. coli and other organisms where advantageous mutations tend to be rare (as an example Elena et al., 1998 and the review from Eyre-Walker et al. 2007)." The authors state that "The proportion of beneficial alleles is not a universal property of the organism. It varies broadly depending on the state of a population (i.e., time of evolution) and external conditions. Hence, the probable reason for the difference is that the cited authors propagate E Coli under different conditions than in 34 whose data we used here."**

**However, since I could not follow the method from the authors and the authors refused to test their method using other reasonable simulations, ...**

How does Reviewer know whether SLIM package is "reasonable", if s/he cannot describe the underlying model and refers to "extensive documentation" instead? Why should we use an obscure program of unknown content, instead of our own codes tested in dozens of our publications in PNAS, Cell, etc?

**...I find it hard to believe this strong claim that goes against a large body of literature showing that advantageous mutations are rare.**

May be, the "body of literature" was not large enough? May be, it was rather SLIM?

**I am unsure why a different environment and time of evolution would cause such a big difference in the amount of advantageous mutations.**

Every biologist knows that an allele deleterious in one environment may be beneficial in another. Examples: drug resistance alleles, immune escape alleles, compensatory alleles, alleles related to melanin production, etc. Suppose a site is uniformly deleterious in the entire population. As time passes, beneficial alleles will gradually accumulate due to natural selection. Eventually, the site will be uniformly beneficial allele. The state in the beginning differs from the state in the end.
