## Supplementary figures and images for "Estimating fitness effects of mutations in the presence of genetic linkage"

### Figure S1

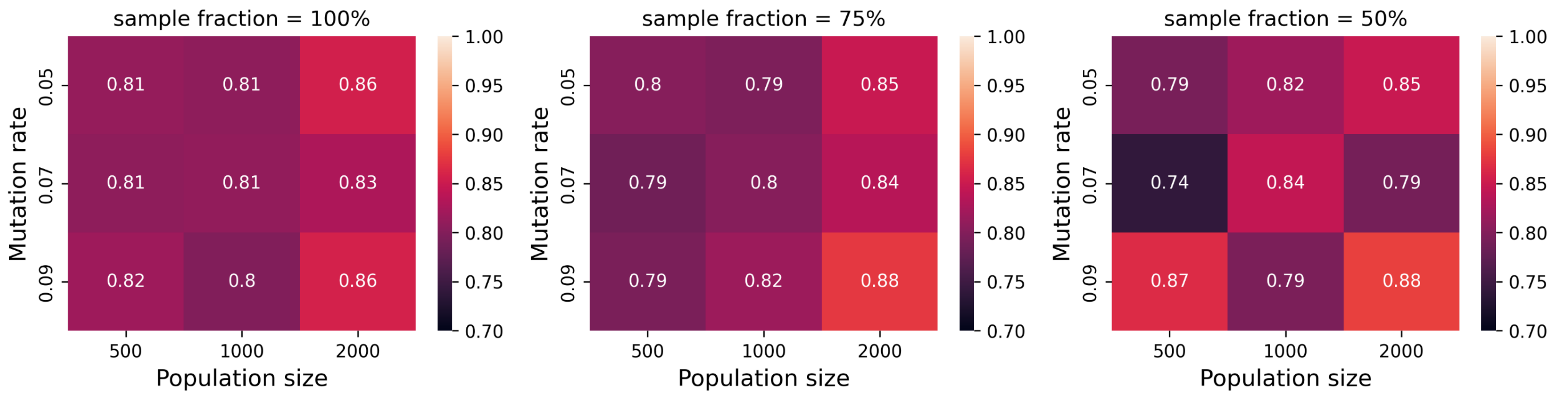
